# Virucidal activity and mechanism of action of cetylpyridinium chloride against SARS-CoV-2

**DOI:** 10.1101/2022.01.27.477964

**Authors:** Nako Okamoto, Akatsuki Saito, Tamaki Okabayashi, Akihiko Komine

## Abstract

**Objective:** Severe acute respiratory syndrome coronavirus 2 (SARS-CoV-2) is the pathogen causing the coronavirus disease 2019 (COVID-19) global pandemic. Recent studies have shown the importance of the throat and salivary glands as sites of virus replication and transmission. The viral host receptor, angiotensin-converting enzyme 2 (ACE2), is broadly enriched in epithelial cells of the salivary glands and oral mucosae. Oral care products containing cetylpyridinium chloride (CPC) as a bactericidal ingredient are known to exhibit antiviral activity against SARS-CoV-2 *in vitro*. However, the exact mechanism of action remains unknown.

**Methods:** This study examined the antiviral activity of CPC against SARS-CoV-2 and its inhibitory effect on the interaction between the viral spike (S) protein and ACE2 using an enzyme-linked immunosorbent assay.

**Results:** CPC (0.05%, 0.1% and 0.3%) effectively inactivated SARS-CoV-2 within the contact times (20 and 60 s) in directions for use of oral care products *in vitro*. The binding ability of both the S protein and ACE2 were reduced by CPC.

**Conclusions:** Our results suggest that CPC inhibits the interaction between S protein and ACE2, and thus, reduces infectivity of SARS-CoV-2 and suppresses viral adsorption.

## 1. Introduction

Coronavirus disease 2019 (COVID-19) has spread worldwide since the end of 2019, and a pandemic was declared by the World Health Organization (WHO) in March 2020. More than 352 million confirmed cases and 5.6 million deaths have been reported as of January 2021 [1]. The novel disease pathogen, severe acute respiratory syndrome coronavirus 2 (SARS-CoV-2), is a single-stranded RNA-enveloped virus. Glycosylated spike (S) proteins cover the surface of SARS-CoV-2, and viral cell entry is mediated by binding to the host cell receptor, angiotensin-converting enzyme 2 (ACE2) [2].

Recent reports have shown that epithelial cells of salivary glands and the throat are major sites of viral replication and release [3,4], as well as infection of gingival epithelial cells in the oral cavity, based on studies of COVID-19 patients [5]. In addition, the Centers for Disease Control and Prevention (CDC) have recommended use of preprocedural oral care products, which may reduce the level of oral microorganisms in aerosols and spatter generated during dental procedures [6]. Many studies have also shown that oral care products containing antiseptic agents decrease the infectivity of SARS-CoV-2 [7-9]. Use of such products may reduce the risk of viral transmission by removal or inactivation of infectious viral particles in the oral cavity [10-14].

Cetylpyridinium chloride (CPC) is a quaternary ammonium compound that is commonly used as a broad-spectrum antimicrobial agent in oral care products. CPC was recently shown to have antiviral activity against SARS-CoV-2 *in vitro* [7,8,15]. The mechanism of action is thought to be viral lipid membrane disruption [16,17], but the effects of CPC on SARS-CoV-2 S protein and ACE2 are unknown. In this study, we evaluated the effects of CPC on reduction of SARS-CoV-2 infection *in vitro* within the contact times for use of oral care products and on the binding ability of S protein and ACE2.

## 2. Materials and Methods

### 2.1 Virus and cells

A strain of SARS-CoV-2 isolated from a patient who developed COVID-19 on the cruise ship Diamond Princess in Japan in February 2020 was obtained from the Kanagawa Prefectural Institute of Public Health (SARS-CoV-2/Hu/DP/Kng/19-027,LC528233). The virus was propagated in VeroE6 cells expressing transmembrane protease serine 2 (TMPRSS2). The cells were obtained from the Japanese Collection of Research Bioresources (JCRB) Cell Bank (https://cellbank.nibiohn.go.jp/english/; JCRB no. JCRB1819. Accessed on 31 May 2021) and cultured in Minimum Essential Medium Eagle with Earle’s salts, L-glutamine and sodium bicarbonate (MEM, pH 7.0-7.6, Sigma-Aldrich, St. Louis, MO, USA) containing 2% fetal bovine serum (FBS, Biowest, Nuaillé, France). At 48 h after infection, virus stocks were collected by centrifuging the culture supernatants of infected cells at 3,000 rpm for 10 min. Clarified supernatants were kept at -80°C until use.

### 2.2 Plaque-forming assay

To evaluate the antiviral effect of CPC on SARS-CoV-2, a plaque-forming assay (PFA) was performed. CPC (Merck KGaA, Darmstadt, Germany) (90 μL) was prepared at a concentration of 0.05, 0.1 or 0.3% (w/v) and mixed with the virus stock solution (10 μL) for 20, 60 or 300 s. Each viral mixture was diluted with Dulbecco’s Modified Eagle Medium (DMEM, Sigma-Aldrich) (×10000) to terminate the CPC reaction. After dilution, the virus solutions were serially diluted in 10-fold steps using serum-free DMEM and then inoculated onto VeroE6/TMPRSS2 cell monolayers in a 12-well plate. After adsorption of virus for 2 h, cells were overlaid with MEM containing 1% carboxymethyl cellulose (Sigma-Aldrich) and 2% FBS (final concentration). Cells were incubated for 72 h in a CO_2_ incubator and then cytopathic effects were observed under a microscope.

Virus suspension mixed with phosphate-buffered saline (PBS(-), Nacalai tesque, Kyoto, Japan) without CPC was used as a negative control. To calculate plaque-forming units (PFUs), cells were fixed with 10% formalin (Fujifilm Wako Pure Chemical, Osaka, Japan) for 30 min, followed by staining with 0.1% methylene blue (Fujifilm Wako Pure Chemical). The antiviral effects of CPC were assessed using the logPFU ratio. Povidone-iodine (PI, final concentration 0.1%) was used as a positive control. All experiments were performed in a BSL-3 laboratory. Three independent experiments were performed for each sample (n = 3).

### 2.3 Degradation analysis of viral protein by Western blot

Western blot analysis was used to determine the effect of CPC on the molecular weight of recombinant SARS-CoV-2 S protein. SARS-CoV-2 Spike Protein (S1+S2 ECD, His tag), Spike S1-His Protein and Spike Protein (RBD, His Tag) were purchased from Sino Biological (Beijing, China). S2-His Protein was purchased from Acro Biosystems (Newark, DE, USA). CPC-treated S proteins were separated by SDS-PAGE and transferred to PVDF membranes. CPC was diluted with distilled water at a concentration of 0.3% (w/v). Each recombinant S protein was diluted with distilled water at a concentration of 444 ng/µL. The protein solution of 6 µL was added to 54 µL of a CPC solution, mixed well, and incubated for 20 s at room temperature. Then, 540 µL of neutralizer (1/10 SCDLP; Fujifilm Wako Pure Chemical) was added to the tube and mixed well. The mixture was diluted 10-fold with 2% FBS containing MEM (Gibco; Thermo Fisher Scientific, Waltham, MA, USA).

For gel electrophoresis, each sample was diluted in 4× Laemmli buffer (Bio-Rad, Hercules, CA, USA) and boiled at 95°C for 5 min. 5 ng of each sample was loaded into a well of a 10% Mini-PROTEAN TGX Precast Gel (Bio-Rad). Gels were transferred to PVDF membranes (Merck KGaA) using a wet-electroblotting chamber system (Bio-Rad) in Towbin buffer (25 mM Tris-HCl [Nacalai tesque], 192 mM glycine [MP Biomedicals, Irvine, CA, USA] and 20% (v/v) methanol [Fujifilm Wako Pure Chemical]). Transfer was performed for 1 h at 4°C. Membranes were washed in PBS (KAC, Kyoto, Japan) containing 0.2% Tween-20 (Sigma-Aldrich) (PBS-T) and blocked with 5% skim milk (Megmilk Snow Brand, Tokyo, Japan) in PBS-T for 30 min. Membranes were incubated overnight at 4°C with a primary antibody for SARS-CoV-2 (2019-nCoV) S protein (Sino Biological, 40589-T62; diluted 1:2,000 in PBS-T). Next, membranes were washed 3 times in PBS-T, incubated with horseradish peroxidase-conjugated anti-rabbit antibody (Abcam, Cambridge, UK) at 1:17,000 dilution for 1 h, washed again 3 times in PBS-T, incubated with SuperSignal West Dura reagent (Thermo Fisher Scientific), and imaged using an Amersham Imager 680 (Cytiva, Tokyo, Japan).

### 2.4 Interaction of the spike protein and ACE2

The interaction between the receptor-binding domain (RBD) of the SARS-CoV-2 S protein and ACE2 was evaluated using a COVID-19 Spike-ACE2 Binding Assay Kit II (RayBiotech Life, Peachtree Corners, GA, USA). The absorbance value (AV) at 450 nm was read using Cytation 5 (BioTek Instruments, Winooski, VT, USA). % binding was calculated as AV of CPC-treated sample / AV of control × 100. Data are presented as the mean ± S.E.M. of triplicate samples of a representative experiment. Similar results were obtained in three independent experiments.

An enzyme-linked immunosorbent assay (ELISA) was performed following the manufacturer’s instructions [18]. Spike RBD protein was mixed with CPC and added to wells coated with ACE2 at final concentrations of CPC of 0.3%, 0.05% and 0.0125%. The mixture in the ACE2-coated wells was incubated for 2.5 h at room temperature. Next, to evaluate the effect of CPC on the RBD, a different assay was performed. Equal amounts of 0.6%, 0.1% or 0.025% CPC solution and RBD solution were mixed, and after 20 s, the mixture was diluted 10-fold with a neutralizer. The solution was then diluted 10-fold with MEM containing 2% FBS and added to the ACE2-coated wells. These samples were then subjected to an ELISA. Finally, another assay was performed to evaluate the effect of CPC on ACE2. 0.3% CPC was added to the ACE2-coated wells and incubated for 20 or 60 s. After removing the CPC solution, neutralizer and MEM containing 2% FBS were added in turn. After removing the liquid in the wells, RBD was added and an ELISA was performed. Assay diluent was used as control. The AV of the sample treated with CPC was compared with the control AV, and the % binding for the sample was calculated.

### 2.5 Statistical analysis

Data are presented as means ± S.E.M. (n=3). Significance was evaluated by a two-sided Dunnett test for comparison with untreated controls. *p < 0.05 and **p < 0.001 indicate a significant difference. IBM SPSS Statistics 27 was used for statistical analysis.

## 3. Results

### 3.1 Virucidal activity of CPC against SARS-CoV-2

To investigate whether CPC has antiviral activity against SARS-CoV-2 to VeroE6/TMPRSS2 cells, a PFA was performed after mixing SARS-CoV-2 with CPC. This revealed that CPC treatment inactivated SARS-CoV-2 within contact times of 20 and 60 s (Table 1). An infectious titer reduction rate of 91.9% was obtained by treatment of virus stock with 0.05% CPC for 20 s. This rate was >97.0% (maximum level of this experiment) with treatment with 0.1% CPC for 20 s, and was also >97.0% with 0.05% CPC treatment for 60 s. These results indicate that CPC has dose- and time-dependent antiviral activity.

**Table 1.**
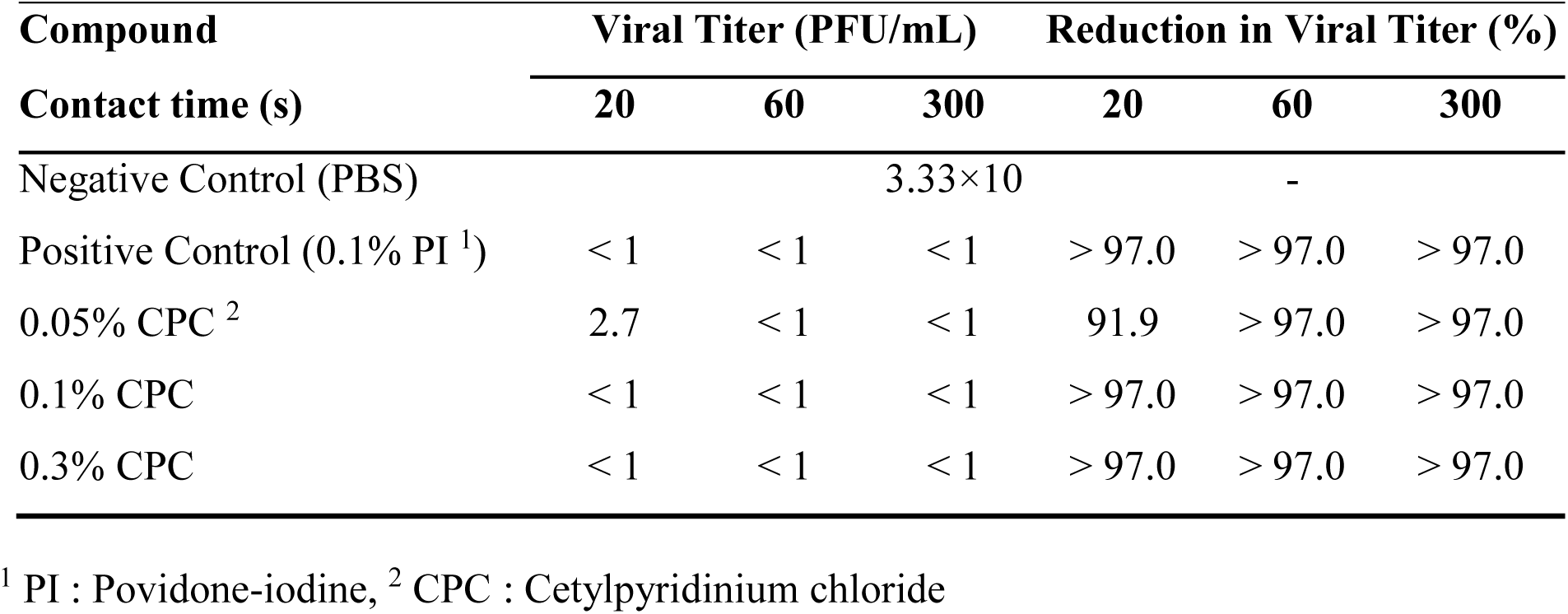
Virucidal activity of CPC against SARS-CoV-2.

### 3.2 Effect of CPC on molecular weight of SARS-CoV-2 S protein

The effect of CPC on SARS-CoV-2 S protein was investigated to examine the virus inactivation mechanism. First, we examined whether CPC caused proteolytic cleavage of the recombinant S protein. In Western blotting after SDS-PAGE under reducing conditions, the molecular weights of four forms of the S protein were unchanged (Fig. 1).

**Figure 1.**
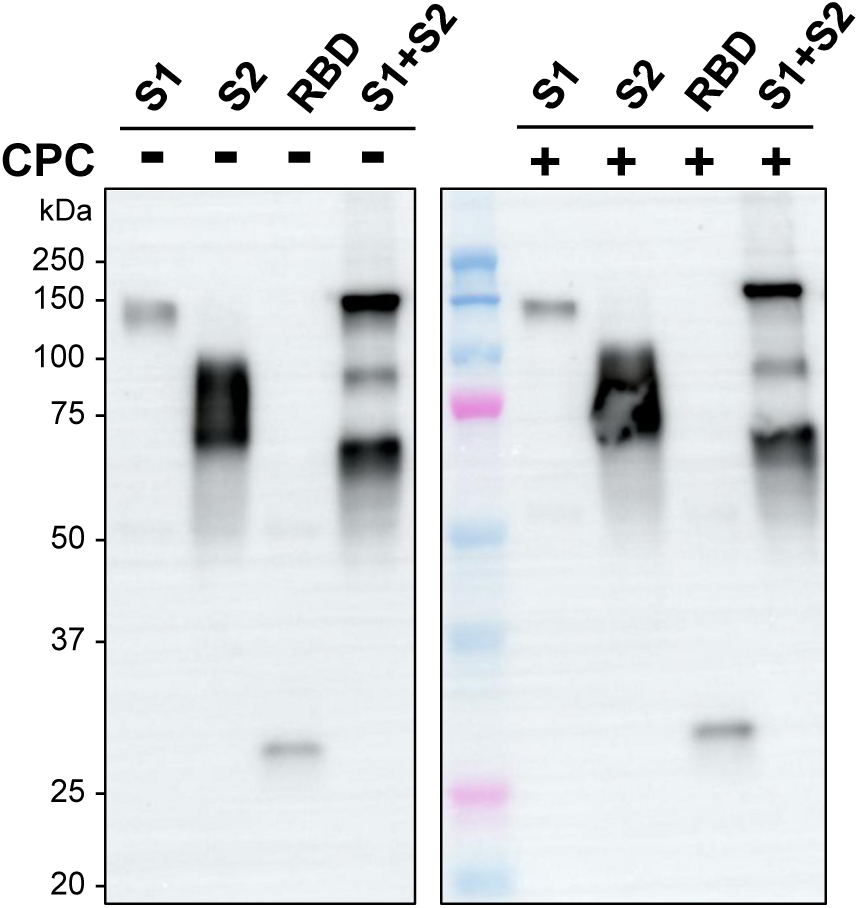
Western blot analysis of CPC-treated SARS-CoV-2 S protein. Four forms of recombinant S protein were separated by SDS-PAGE after treatment with 0.3% CPC for 20 s and neutralization. All samples and protein standards were run on the same blot.

### 3.3 Interaction assay between spike RBD and ACE2

To evaluate the effect of CPC on the activity of S protein, the interaction between the RBD of S protein and ACE2 was analyzed by ELISA in three different assays. First, the RBD and ACE2 were incubated for 2.5 h in the presence of CPC. The % binding rate was significantly reduced with 0.3% and 0.05% CPC (Fig. 2A). Next, a RBD-ACE2 binding assay after CPC treatment of the RBD and neutralization showed that % binding was significantly reduced with 0.3% and 0.05% CPC (Fig. 2B). In this assay, the influence of neutralized CPC solution on binding of ACE2 was also investigated (Fig. 2B: “Interference”). Under these conditions, binding was equivalent to that with 0% CPC (control), indicating no interference of neutralized CPC solution on binding of ACE2. Finally, a RBD-ACE2 binding assay after CPC treatment of ACE2 and neutralization showed that the % binding of CPC-treated ACE2 decreased by about 70% compared to the untreated control (Fig. 2C).

**Figure 2.**
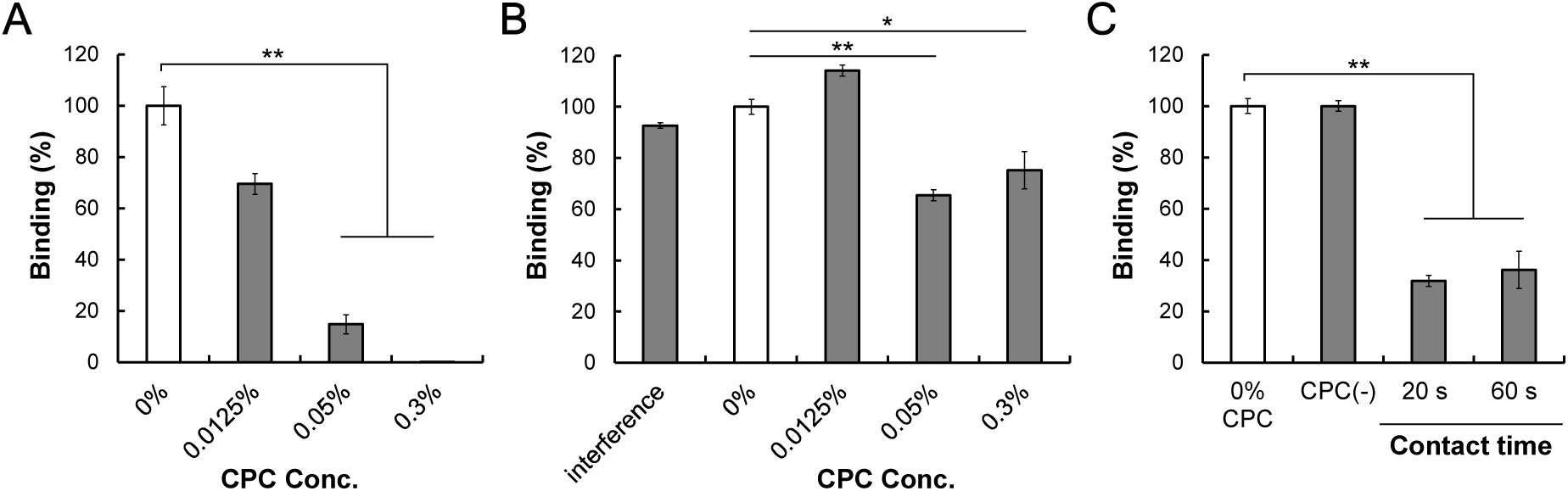
Effects of CPC on the interaction between S protein (RBD) and ACE2. (A) Binding in the presence of CPC. The % binding of each sample was calculated based on the absorbance with 0% CPC (control). (B) Binding of CPC-treated RBD with ACE2. “Interference” indicates the influence of a neutralized CPC solution on binding of RBD to ACE2. After neutralization, CPC was incubated in ACE2-coated wells for 1 h and the ACE2-RBD binding assay was performed. (C) Binding of RBD with 0.3% CPC-treated ACE2. The assay diluent was tested similarly and is indicated as CPC(-). Data are shown as means ± S.E.M. (n=3). *p < 0.05, **p < 0.001 vs. untreated control by two-sided Dunnett test.

## 4. Discussion

This study showed that CPC reduces the infectivity of SARS-CoV-2 below the detection limit at a concentration of CPC above 0.05% in treatment within 60 s. CPC inhibited the interaction between S protein and ACE2 without cleavage of S protein. These results indicate that CPC has dose- and time-dependent antiviral activity. Viral envelope disruption by CPC has been proposed as a mechanism of inactivation for an enveloped-virus [19] and this may also occur in SARS-CoV-2 inactivation [16,17]. In the current study we considered other mechanisms through examining the effect of CPC on the viral S protein. Western blot analysis showed that cleavage of S protein was not induced by CPC, and an ELISA showed reduced binding of S protein to ACE2. The results suggest that CPC may influence the S protein through conformational change, rather than proteolytic cleavage.

An evaluation of the interaction of CPC with human serum albumin (HSA) using isothermal titration calorimetry indicated that HSA was unfolded in two steps depending on the CPC concentration, resulting in loss of secondary and tertiary structures [20]. In the current study, the CPC-treated S protein showed reduced binding to ACE2 without proteolytic cleavage, which suggests that the mechanism of action of CPC involves denaturation of S protein. Thus, the antiviral activity of CPC against SARS-CoV-2 may be attributable to this mechanism. Our results also suggest that CPC may suppress the activity of human ACE2 and reduce the efficiency of infection of host cells without proteolytic cleavage.

The results for binding of CPC-treated RBD and ACE2 (Fig. 2B) indicated that the effect of CPC on binding was not dose-dependent. These findings may be due to a change in micelle morphology in aqueous solution that depends on the CPC concentration. CPC is a cationic surfactant with strong antibacterial and antifungal activity [21], and tends to form micelles by association via hydrophobic chains at high aqueous concentrations above the critical micelle concentration (CMC). The mechanism of protein denaturation by ionic surfactants involves binding of a monomeric surfactant to specific sites of the protein via electrostatic and hydrophobic interactions, rather than binding as micelles [22]. The amount of monomers binding to the protein is constant even at free surfactant concentrations beyond the CMC, and thus, the rate of unfolding tends to level off around the CMC [22]. Given that the CMC of aqueous CPC solution is 0.9-1.0 mM [23] (approximately 0.03% w/v) at 25°C, the number of CPC monomers that bind to proteins is about constant at a concentration above 0.03%.Therefore, the absence of dose-dependent binding inhibition is due to the high concentration of CPC above its CMC, at which binding of monomers was saturated and the effect of CPC reached a plateau.

## 5. Conclusions

CPC exhibited virucidal activity against SARS-CoV-2 at a concentration above 0.05%. One mechanism of this phenomenon may be related to denaturation of the viral S protein (RBD) and reduction in binding to the host ACE2 receptor. Our results also suggest that CPC has an influence on human ACE2 and suppresses viral adsorption.

## Acknowledgments

None

